# The Insulin Receptor in Astrocytes is Involved in the Entrance of Circulating Insulin into the Brain

**DOI:** 10.1101/720813

**Authors:** Ana M. Fernandez, M. Navarrete, Jose Carlos Davila, Cristina Garcia-Caceres, Rocio Palenzuela, Samuel Ruiz de Martin Esteban, Ricardo Mostany, Matthias Tschöp, Antonia Gutierrez, Ignacio Torres Aleman

## Abstract

Circulating insulin enters into the brain through mechanisms that are not yet entirely understood. We now report that mice lacking insulin receptors (IR) in astrocytes show reduced uptake of circulating insulin and blunted brain responses to this circulating hormone, suggesting that astrocytes form part of the cellular pathway in the entrance of circulating insulin into the brain. Since we previously showed that ablation of IR in astrocytes decreases brain glucose uptake, we conclude that astrocytic IRs regulate uptake of circulating glucose and insulin.

## Introduction

The origin of insulin in the brain has been a matter of debate until relatively recently (Gray et al., 2014). Early observations documented the presence of insulin in the cerebrospinal fluid (CSF) and the brain (Margolis and Altszuler, 1967; Pansky and Hatfield, 1978). Because expression of insulin by brain cells was firmly established only recently (Kuwabara, 2011), for many years, insulin actions in the brain were assumed to arise from pancreatic insulin (Schwartz and Porte, 2005) that has to cross the blood-brain-barriers (BBBs). Indeed, there are insulin receptors (IR) in the cells forming the BBBs; that is, epithelial cells at the choroid plexus (Baskin et al., 1986), brain capillary endothelial cells (Frank et al., 1986) that transcytose insulin (King and Johnson, 1985), and astrocytes (Baron-Van et al., 1991). Accordingly, previous observations indicated that insulin from the circulation enters the brain through a transport mechanism involving its receptor in brain endothelial cells forming the BBB (Gray et al., 2017), although more recent observations indicate that the process may be IR independent (Hersom et al., 2018; Rhea et al., 2018).

Astrocytes seal brain endothelial cells of brain capillaries by wrapping them with their end-feet (Mathiisen et al., 2010). In this way, astrocytes constitute an additional cell layer of the BBB (Zlokovic, 2008). Once insulin has being transported across endothelial cells (King and Johnson, 1985), it may be taken up by astrocytes either to be further transported into the brain parenchyma through transcytosis (Simionescu et al., 2002), to modulate the activity of these glial cells (Cai et al., 2018), or both. To help clarify the route of entrance of circulating insulin into the brain, we analyzed its brain uptake in mice lacking IR in astrocytes. Ablation of IR in astrocytes blocked the entrance of circulating insulin into the brain, suggesting that astrocytes form part of the intracellular pathway of passage of this hormone into the brain.

## Results

### IRs are present in astroglial end-feet

We first determined whether IR are present in astrocytic end-feet ensheathing endothelial cells of the BBB, as its presence in this compartment will allow circulating insulin to readily access astrocytes. Using combined IR immunogold labeling and electron microscopy we localized these receptors in astrocytic end-feet (Figure 1A,B). This anatomical location, in close proximity to endothelial cells, together with the expression of IR in the latter type of cells (Figure 1B), or as an yet uncharacterized transporter (Rhea et al., 2018), would allow the passage of circulating insulin into astrocytes by transcytosis through endothelial cells (Gray et al., 2017).

**Figure 1:**
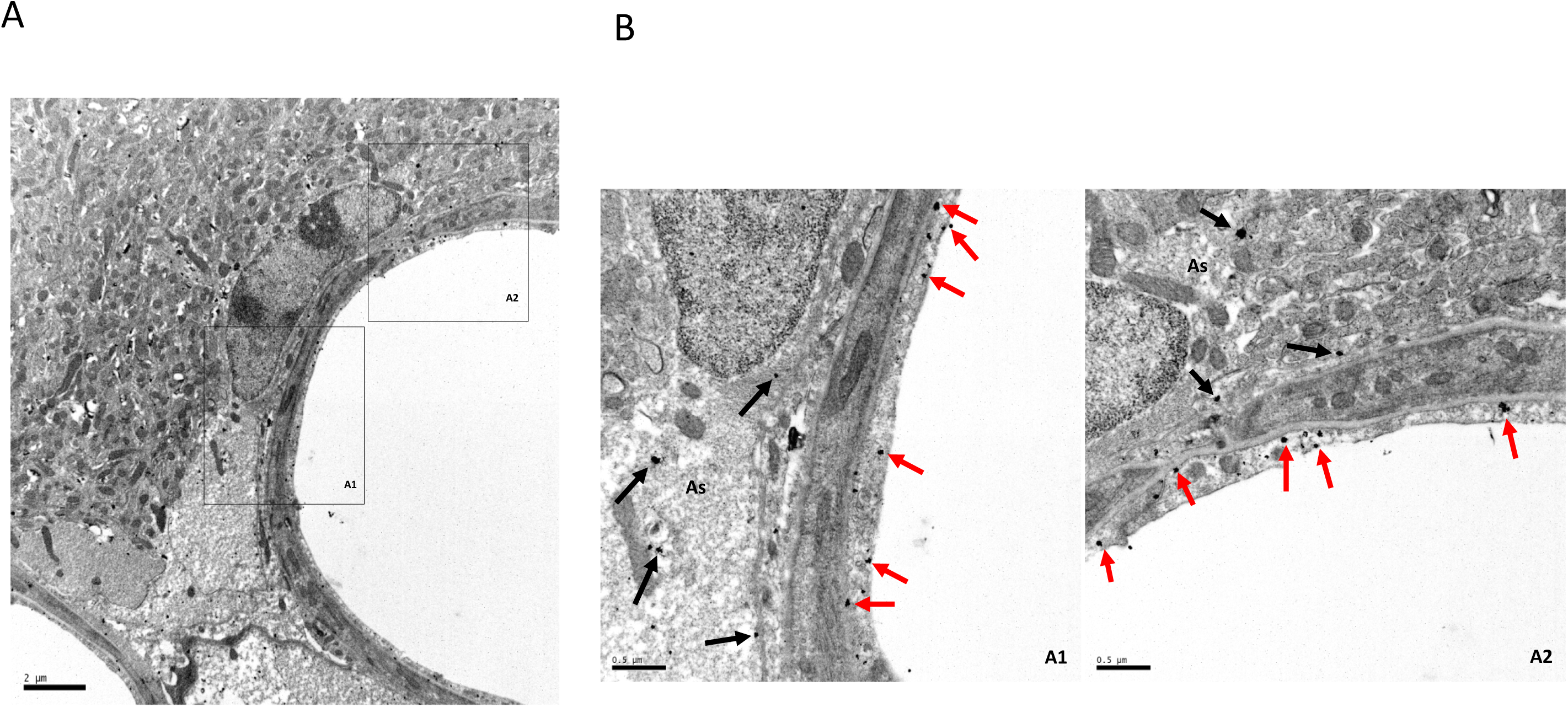
IR is expressed by astrocyte end-feet. **A**, Electron microscopy photograph using immuno-gold labeling for IR illustrating the presence of immunoeractive IR deposits in astroglial end-feet surrounding brain capillaries and in endothelial cells. **B**, A1 and A2 squares from A, are shown at greater magnification. Note the presence of IR gold particles in endothelial cells (red arrows) and astrocytic end-feet (black arrows). Bar in A is 2 µm, and in B, 0.5 µm.

### Astrocytic IRs are required for brain IR activation and stimulation of astrocyte signaling by circulating insulin

As already documented (Sartorius et al., 2015), systemic administration of insulin results in robust activation of brain IR (Figure 2A). However, GFAP IR KO mice show attenuated responses to peripheral insulin injection, as determined by reduced phosphorylation of brain (Figure 2B), but not muscle IR (Figure 2C). The reduction in brain IR phosphorylation was not due to lower brain IR levels in mutant mice as a result of the ablation of astrocytic IR since receptor activation was normalized by its relative levels (similar between tamoxifen-treated and non-treated mice; see lower blot in Figure 2B). Furthermore, GFAP IR KO mice show normal brain insulin-like growth factor I (IGF-I) receptor phosphorylation in response to systemic injection of this close relative of insulin (Suppl Fig 1A), indicating specific loss of sensitivity to systemic insulin.

Since changes in intracellular Ca^2+^ levels are used as a proxy of astrocyte activity (Ma et al., 2016), we next determined whether circulating insulin modulates intracellular Ca^2+^ in astrocytes of GFAP IR KO mice. We monitored in vivo Ca^2+^ levels in astrocytes located in the somatosensory cortex by two-photon laser-scanning fluorescence microscopy using a genetically-encoded Ca^2+^ indicator (Lck-GCaMP6f) (Figure 3A,B). We found that systemic administration of insulin (3 IU/kg) elicited Ca^2+^ spikes frequency increases in control littermates (Figure 3C-D). In contrast, these calcium activity increases were virtually absent in transgenic mice lacking insulin receptors in astrocytes (GFAP IR KOs), confirming that astrocytic IRs are necessary for astrocytes to respond to systemic insulin.

**Figure 2:**
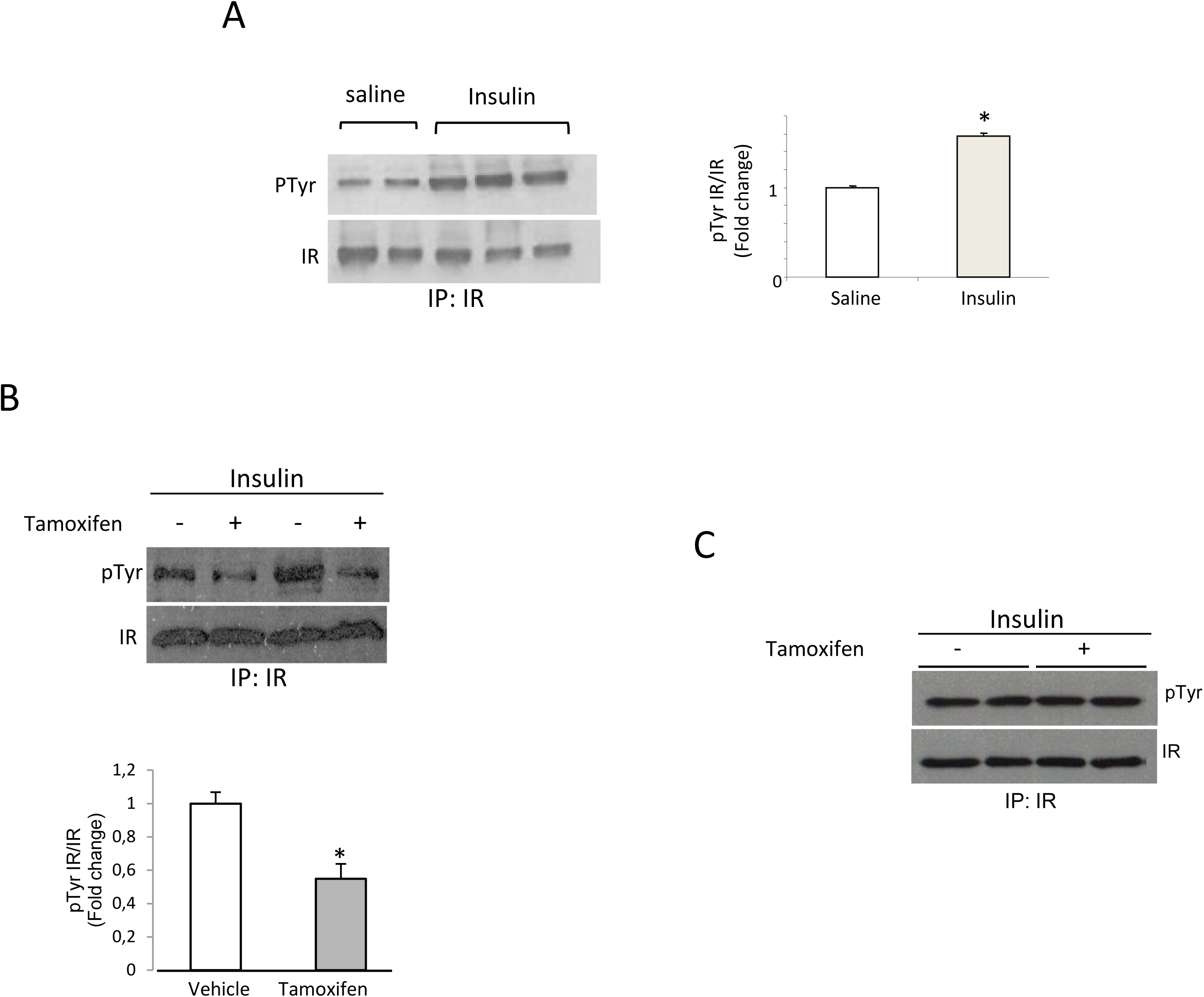
Systemic injection of insulin stimulates IR phosphorylation in cortex. **A**, Intraperitoneal injection of insulin (5 U/ml) results in stimulation of IR in the mouse somatosensory cortex as compared to control mice (injected with the same amount of saline). **B**, Ablation of IR in GFAP IR KO astrocytes after tamoxifen administration results in markedly reduced activation of cortical IR after systemic insulin in comparison to vehicle (corn oil) injected mice. **C**, In these animals, activation of muscle IR was intact, as compared to oil-injected GFAP IR KO mice. Activation of IR is shown as amount of Tyrosine phosphorylated receptor (pTyr) normalized by total levels of immunoprecipitated IR. Representative blots are shown. Quantification of changes is shown in histograms (n=8 for each group; *p<0.05 vs respective control mice).

### Astrocytic IRs are involved in brain uptake of circulating insulin

To analyze the pathways involved in the passage of circulating insulin into the brain with greater detail, we injected mice with digoxigenin-labelled insulin (Dig-Ins) in the carotid artery and examined its route of entrance by immunolabelling. In control GFAP IR KO mice (not injected with tamoxifen) neuronal digoxigenin staining was readily seen (Figure 4A, panels b-c, and B). However, in tamoxifen-treated GFAP IR KO mice, neuronal staining was lost (Figure 4C, panels b-c, and D). In addition, a strong Dig-Ins staining in vessels was seen in these animals (Figure 4C, panel a, and Suppl Fig 1B), suggesting that it is retained inside the endothelium without crossing into the brain parenchyma.

**Figure 3:**
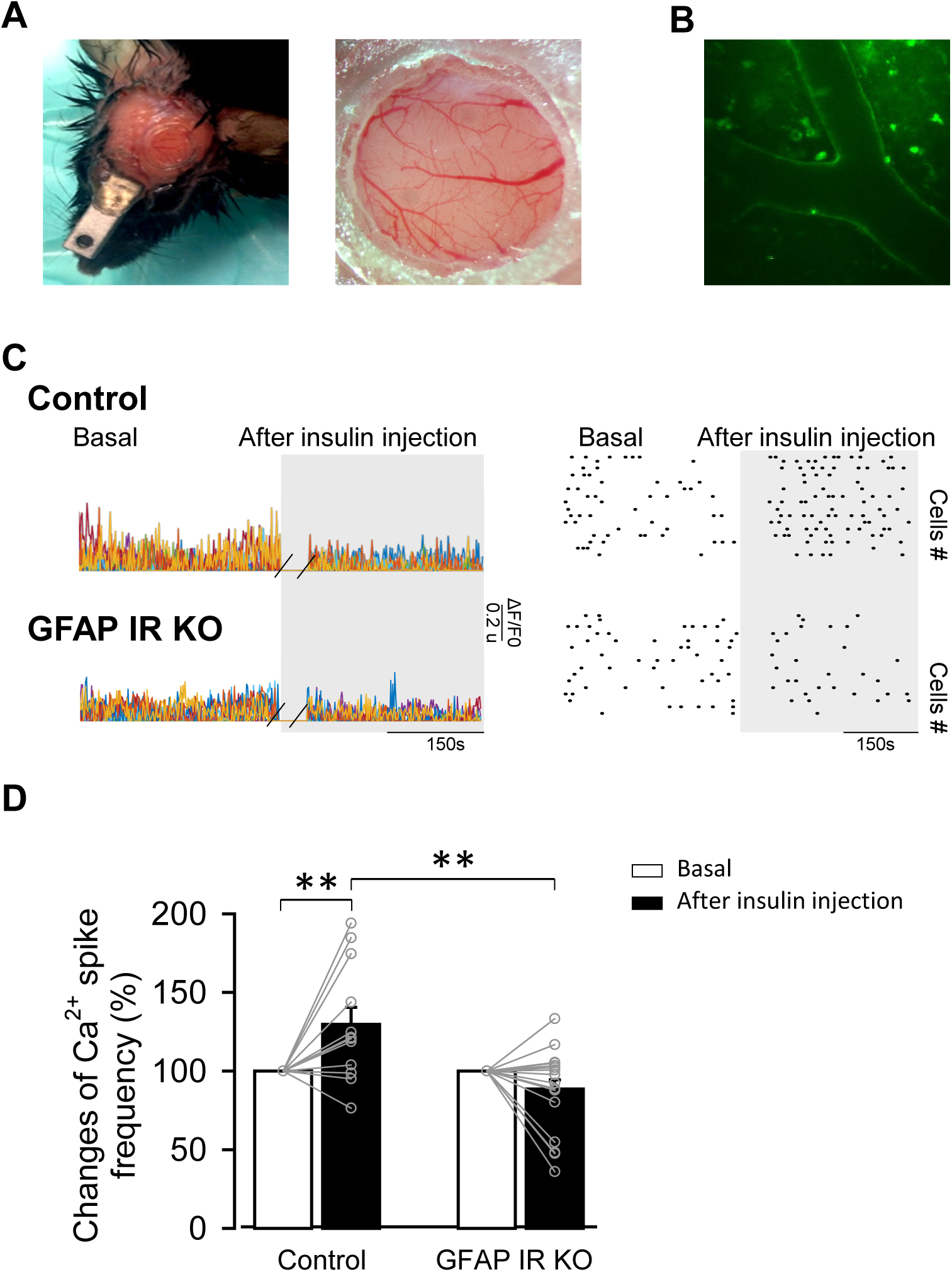
In vivo two-photon imaging of activity of GCaMP6-labeled astrocytes. **A**, Mouse bearing a cranial window (left), and a macroscopic image of the mouse cranial window (right). **B**, Representative two-photon fluorescence intensities for viral transfection of AAV5-P_GFAP_-Lck-GCaMP6f in somatosensory cortex imaging *in vivo*. Note astrocytic endfeet enwrapping blood vessels. Scale bar, 35 mm. **C**, Left, Representative experiments showing the amplitude of calcium events versus time 5 min before (basal) and 15 min after ip injection of insulin (3 IU/kg body weight) in Control littermates (top), and GFAP IR KO (bottom) mice. Right, Representative raster plot of ROIs activity versus time showing the frequency of calcium events 5 min before (basal) and 15 min after injection of insulin in Control (top) and GFAP IR KO (bottom) mice. D, Changes of spike frequency of Ca^2+^ signals per area 5 min before (basal) and 15 min after i.p. injection of insulin in Control (n = 12 from n = 4 mice, P < 0.01 Wilcoxon test) and GFAP IR KO (n = 19 from n = 4 mice, P = 0.163 Wilcoxon test) mice. Differences between groups were determined by Kruskal-Wallis test (** P < 0.01).

**Figure 4:**
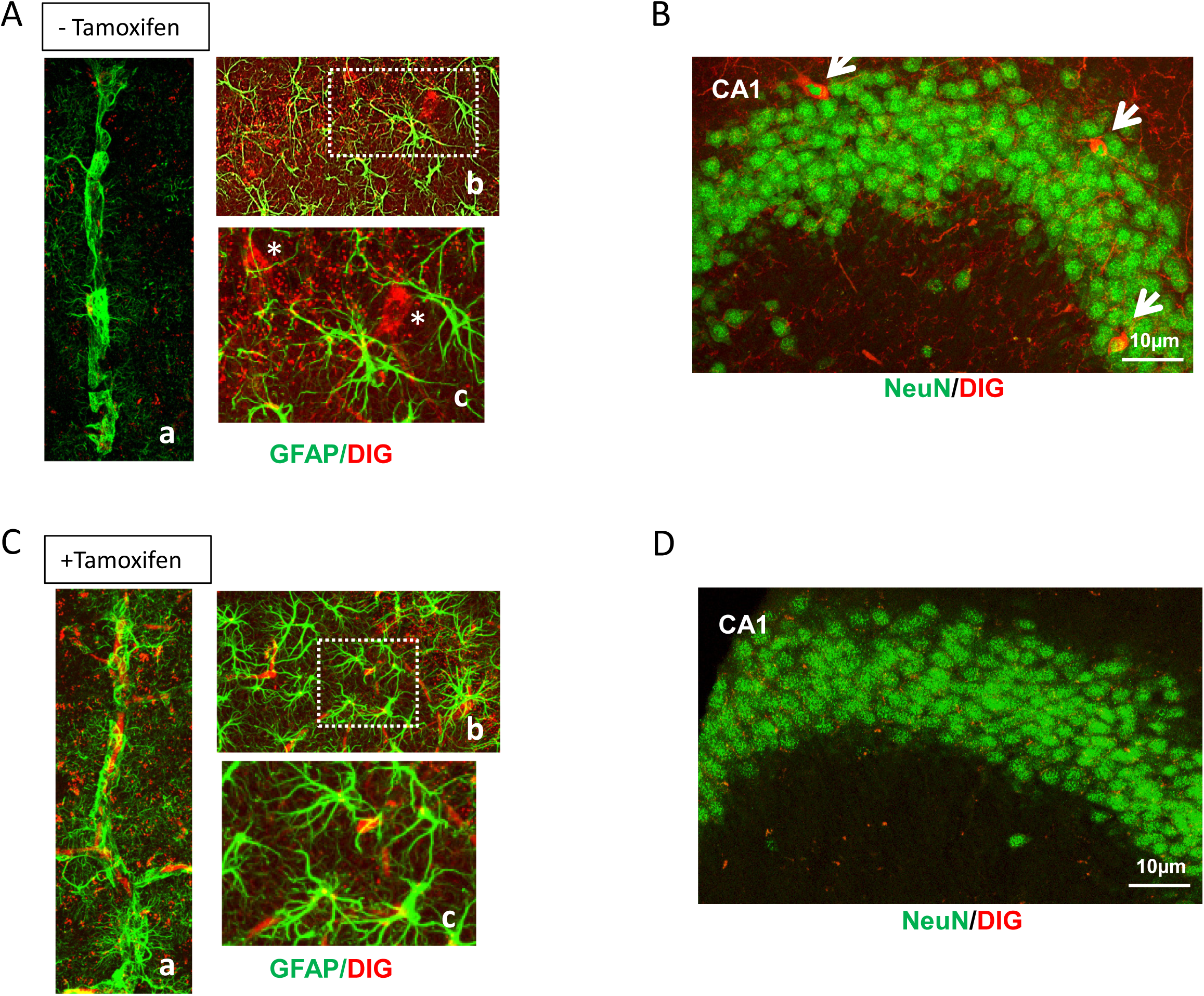
Insulin receptors in astrocytes are required for neurons to capture circulating insulin. **A**, Intra-carotid injection of digoxigenin-labeled insulin (Dig-insulin) results in its neuronal accumulation only when astrocytes express IR (no tamoxifen). Panel a shows astrocyte end-feet (GFAP^+^ in green) ensheathing brain capillaries in control (-tamoxifen) mice receiving Dig-insulin injections. No Dig staining is seen. In contrast, Dig staining is seen in cell of panel b (white dotted square) with neuronal morphology (red), that are magnified in panel c (white asterisks). **B**, Accumulation of labeled insulin is readily seen in hippocampal neurons (NeuN^+^ cells) of control mice (no tamoxifen). **C**, GFAP IR KO mice (treated with tamoxifen) do not show Dig staining in neurons after intracarotid injection of Dig-insulin. In panel a, vessels surrounded by GFAP^+^ astroglial end-feet show abundant digoxigenin staining, suggesting lower transfer of Dig-insulin into the brain. In panels b and c (higher magnification of dotted white square in b) neuronal-like shapes with Dig staining are absent. Note the pronounced GFAP reactivity of astrocytes as compared to panel c in A. **D**, GFAP IR KO (+tamoxifen) do not show NeuN^+^ (green) cells co-labelled with Dig in the hippocampal granular cell layer. Green: GFAP^+^ or NeuN^+^ cells; Red: Digoxigenin^+^ cells.

## Discussion

The first cell layer separating the brain parenchyma from the circulation through the BBB is formed by brain endothelial cells. Whether insulin receptors in these cells are involved in the transport of circulating insulin into the brain, is controversial (King and Johnson, 1985; Gray et al., 2017; Hersom et al., 2018; Rhea et al., 2018). The second cell layer of the BBB is formed by astrocytic end-feet. The present observations indicate that insulin receptors in astrocytes are involved in the entrance of circulating insulin into the brain. Moreover, these receptors are also involved in astrocyte activation by circulating insulin. Once insulin is transcytosed by brain endothelial cells through a regulated process, involving or not its receptor (Banks, 2004; Gray et al., 2017), it is then taken up by astrocytes in a receptor-mediated process to deliver it to neighboring neurons. Hence, circulating insulin access astrocytes not only to modulate their function (Garcia-Caceres et al., 2016; Cai et al., 2018) and activity (present results), but also as a gate of entrance to neurons, as blockade of this pathway results in blockade of neuronal uptake.

Although circulating insulin plays many important roles in the brain, the key question of how it reaches its numerous brain targets remained, until now, poorly characterized. Indeed, it might be of great relevance to know the route of entrance of serum insulin into the brain as in diabetes and several neurodegenerative diseases, brain insulin resistance is a likely pathogenic component. Knowing whether reduced entrance of insulin into the brain and/or loss of sensitivity of brain cells underlie reduced brain insulin function in these pathologies will greatly assist new therapeutic options. In conjunction with recent observations that mice without IR in astrocytes show a depressive phenotype (Cai et al., 2018), the present results pose astrocytic IRs as important targets for future therapies.

## Materials and Methods

### Animals

Wild type 6 month-old male mice (C57BL6/J) were used in EM experiments. Also, mice with insulin receptors ablated in astrocytes (GFAP IR KO mice) under regulation of tamoxifen were generated as described in detail before (Garcia-Caceres et al., 2016), by crossing hGFAP-CreER^T2^ on a C57 BL/6J background (FM Vaccarino, Yale Univ) with IR^f/f^ mice having the IR gene flanked by loxP sites (R Kahn, Joslin Diabetes Center). Intraperitoneal injection of tamoxifen for 5 consecutive days (75 mg/kg) to adult GFAP IR KO mice (4-5 weeks old) eliminated IR specifically in astrocytes, as reported (Garcia-Caceres et al., 2016), and resulted in significantly decreased IR content in brain (Suppl Fig 1C). Controls received vehicle injections (corn oil). GFAP IR KO mice were genotyped using PCR protocols already detailed (Garcia-Caceres et al., 2016). Mice had access to food and water *ad libitum* and were kept under light/dark conditions. Animal procedures followed European (86/609/EEC b 2003/65/EC, European Council Directives) guidelines and studies were approved by the respective local Bioethics Committees (CAM PROEX 01/14).

### Reagents

The following reagents were used: tamoxifen (Sigma, Darmstadt, Germany), Insulin (Sigma), IGF-1 (PreproTech, Rocky Hill, NJ, USA), and tomato lectin-FITC (Sigma),. Antibodies used were: Phosphotyrosine (clone PY20, BD Transduction laboratories, San Jose, CA, USA), GFAP (Dako, Glostrup, DK), NeuN (Millipore, Darmstad, Germany), Digoxigenin (Roche Diagnostics, Mannheim, Germany), Insulin receptor ß (Santa Cruz Biotechnology, Dallas, TX, USA), IGF-1 receptor ß (Cell Signaling Technology, Danvers, MA, USA) and ß-actin (Sigma,).

### Digoxigenin-labeled insulin

100 µl of 1 mg/ml recombinant human insulin (Sigma) was mixed with 1 ml of PBS pH 8.5. Digoxigenin-3-O-methylcarbonyl-έ-aminocaproic acid-N-hydroxysuccinimide ester (NHS-Dig; Roche Diagnostics) dissolved in dimethyl sulfoxide (DMSO) was added to the insulin solution at a molar ratio of 1:5. After incubation at room temperature for 2 h, non-reacted NHS-Dig was removed by a Sephadex G-25 column (GE Healthcare, Chicago, IL, USA). Digoxigenin-labeled insulin (Dig-insulin) was stored at −20°C until use.

### Insulin administration

Mice were anesthetized with isofluorane (with oxygen flux at 0.8–1 l/min). The common carotid artery was exposed and the external carotid ligated. A guide cannula was inserted in the common carotid artery and 100µl Dig-insulin was injected using a Hamilton syringe coupled to a nanoinjector. After 1 hour, mice were perfused with saline and 4% paraformaldehyde. Brains were removed and processed for immunohistochemistry. For IR phosphorylation experiments, mice were injected with 5 IU insulin intraperitoneally. After 1h, mice were perfused with saline and brain and skeletal muscle isolated, frozen until assayed by IR immunoprecipitation and Western blotted with anti-pTyr.

### Cranial window surgery for *in vivo* imaging

Chronic glass-covered cranial windows were implanted at least 2 weeks before the beginning of the in vivo calcium imaging of astrocytes (Figure 3A). Briefly, mice were anesthetized (isoflurane, 5% for induction, 1.5% for maintenance via nose cone) and placed on a stereotaxic frame. Dexamethasone (0.2 mg/kg) and carprofen (5 mg/kg) were administered. A 4 mm-diameter craniotomy was performed with a pneumatic dental drill over the somatosensory cortex. A stereotaxic microinjection (400 nl; 30 nl/min) of AAV2/5-P_GFAP_-Lck-GCaMP6f (PENN Vector Core; viral titer 6.13 x 10^13^) was made. A round glass coverslip (5 mm) was laid over the dura mater, covering the exposed brain and part of the skull and glued to the latter with cyanoacrylate-based glue. A layer of dental acrylic was then applied throughout the skull surface and up to the edges of the coverslip. A titanium bar (9.5 × 3.2 × 1.1 mm) was embedded in the dental acrylic to secure the mouse onto the stage of the microscope for imaging.

### Ca^2+^ imaging

Imaging of GCaMP6f-expressing astrocytes was performed with a two-photon laser scanning microscope, custom-modified with a femtosecond laser (Chameleon Ultra II, Coherent Inc) and ScanImage 3.8 software written in MATLAB (MathWorks; RRID:SCR_001622), under isoflurane anesthesia (1%−1.5%). Mice were secured to the microscope using the titanium head bar. A 10X immersion objective (numerical aperture, NA, of 0.45, Nikon) was used to create a map of the area of interest and a 40X water-immersion objective (NA=1.1) was used for visualization of individual areas. Excitation of GFP was achieved by tuning the laser at 780 nm with a power at the sample of < 20mW. After two weeks of surgery and viral injection, specific expression of constructs in the astrocytes was confirmed by immunostaining. Images were acquired every 1.4-1.7 seconds and ROIs were designed with ImageJ in a semi-automated manner using the GECIquant program. Ca^2+^ variations were estimated as changes of the fluorescence signal over baseline (ΔF/F_0_), and regions of interest were considered to respond to the stimulation when ΔF/F_0_ increased three times the standard deviation of the baseline. The astrocyte Ca^2+^ signal was quantified from the Ca^2+^ oscillation frequency. The time of occurrence was considered at the onset of the Ca^2+^ spike. Custom-written software in MATLAB (MATLAB R2016; Mathworks, Natick, MA) was used for computation of fluorescence of each ROI.

### Immunofluorescence

Immunolabeling was performed as described (Fernandez et al., 2012). Mice were deeply anesthetized with pentobarbital (50 mg/kg), and perfused transcardially with saline followed by 4% paraformaldehyde in 0.1 M phosphate buffer, pH 7.4 (PB). Brains were dissected, and coronal 50-µm-thick brain sections were cut in a vibratome and collected in PB. Sections were incubated to block non-specific antibody binding, followed by incubation overnight at 4°C with the primary antibodies in PB – 1% bovine serum albumin – 1% Triton X-100 (PBT). After several washes in PBT, sections were incubated with the corresponding Alexa-coupled secondary antibody (1:1000, Molecular Probes, ThermoFisher Scientific) diluted in PBT. Again, three 5-minute washes in PBT were required. Slices were rinsed several times in PB, mounted with gerbatol mounting medium, and allowed to dry. For endothelial cell staining, sections were incubated with Tomato lectin-FITC overnight at 4°C followed by several washes in PBT. Omission of primary antibodies was used as control. Confocal analysis was performed in a Leica microscope (Wetzlar, Germany).

### Immunoprecipitation and Western blotting

Immunoassays were performed as described (Fernandez et al., 2007). Brain tissue was homogenized in ice-cold buffer with 10mM Tris HCl pH 7.5, 150mM NaCl, 1mM EDTA, 1mM EGTA, 1% Triton X-100, 0.5% NP40, 1mM sodium orthovanadate, and a protease inhibitor cocktail (Sigma) plus 2mM PMSF, using 1 ml of buffer per mg of tissue. Insoluble material was removed by centrifugation, and supernatants were incubated overnight at 4 °C with the antibodies. Immunocomplexes were collected with Protein A/G agarose (Santa Cruz Biotechnology) for 1 h at 4 °C and washed 3X in homogenization buffer before separation by SDS-polyacrylamide gel electrophoresis and transferred to nitrocellulose membranes. After blocking for 1 h with 5% BSA in TTBS (20 mM Tris-HCl, pH 7.4, 150 M NaCl, 0.1% Tween 20), membranes were incubated overnight at 4 °C with the different antibodies in TTBS, washed, incubated with secondary antibodies and develop using the Odissey procedure (Li-Cor Biosciences, Lincoln, NE, USA).

### Immunoelectron microscopy

Immunogold procedure was performed as previously described (Gomez-Arboledas et al., 2018). After deep anesthesia with sodium pentobarbital (60 mg/kg), 6 month-old male mice (C57BL6/J) were perfused transcardially with 0.1 M phosphate-buffered saline (PBS), pH 7.4 followed by fixative solution containing 4% paraformaldehyde, 75 mM lysine and10 mM sodium metaperiodate in 0.1 M PB, pH 7.4. Brains were removed, post-fixed overnight in the same fixative solution at 4°C, coronally sectioned at 50 μm thicknesses on a vibratome (Leica VT1000S), and serially collected in wells containing cold PB and 0.02% sodium azide. For IGF-1R or IR immunogold labelling, sections containing the somatosensory cortex were used. Sections were first washed with PBS and incubated in a 50 mM glycine solution 5 minutes to increase antibody binding efficiency. Following a standard immunohistochemical protocol, tissue was first free-floating incubated in a rabbit polyclonal anti-IGF-IRα antibody or a anti-IR antibody in a PBS 0.1M/1% BSA solution for 48 hours at 22°C. Then, sections were washed in PBS, and incubated with 1.4 nm gold-conjugated goat anti-rabbit IgG (1:100; Nanoprobes) overnight at 22°C. After post-fixing with 2% glutaraldehyde and washing with 50 mM sodium citrate, labelling was enhanced with the HQ Silver™ Kit (Nanoprobes), and gold toned. Finally, immunogold labelled sections were fixed in 1% osmium tetroxide, block stained with uranyl acetate, dehydrated in acetone, and flat embedded in Araldite 502 (EMS, USA). Selected areas were cut in ultrathin sections (70-80 nm) and examined and photographed with a JEOL JEM1400 electron microscope. As a control for the immunogold technique, sections were processed as above but omitting the primary antibody. No specific labelling was observed in these control sections.

### Statistics

Statistical analysis was performed using GraphPad Prism 5.0 (La Jolla, CA, USA). To compare differences between two groups, Student’s t-test (Gaussian distribution of data) or Mann–Whitney test (non-Gaussian distribution of data) were used. To compare multiple variables, two-way ANOVA was used, followed by a Bonferroni post hoc test to compare replicate means. Statistical differences were considered when p < 0.05. Results are presented as mean ± SEM of five to eight mice per group. (*p< 0.05, **p< 0.01, ***p< 0.001).

## Acknowledgments

We are thankful to M. Garcia for technical support. This work was funded by a grant from Ciberned, an Inter-CIBER project (PIE14/00061), and SAF2016-76462-C2-1-P (MINECO) to ITA, and grant PI18/01557 (to AG) from Instituto de Salud Carlos III (ISCiii) of Spain (co-financed by FEDER funds from European Union).

## LEGENDS TO FIGURES

**Supplementary Figure 1**: **A**, Systemic injection of IGF-I to GFAP IR KO mice stimulates phosphorylation of its receptors in cortex. **B**, Digoxigenin staining in brain vessels (tomato lectin^+^ cells) of control (no tamoxifen) and GFAP IR KO mice (+ tamoxifen). **C**, GFAP IR KO mice show reduced IR levels in brain, as compared to controls (treated with tamoxifen vehicle; **p<0.01 vs control).

## Notes

#### Summary of Updates

We added new experiments that reinforce our conclussions. These appear as new Figure 3 with the corresponding new text in results and discussion. New authors have been added since they performed the new experiments.

