## Supplementary material for "The Insulin Receptor in Astrocytes is Involved in the Entrance of Circulating Insulin into the Brain": Suppl Figure

### Slide 1
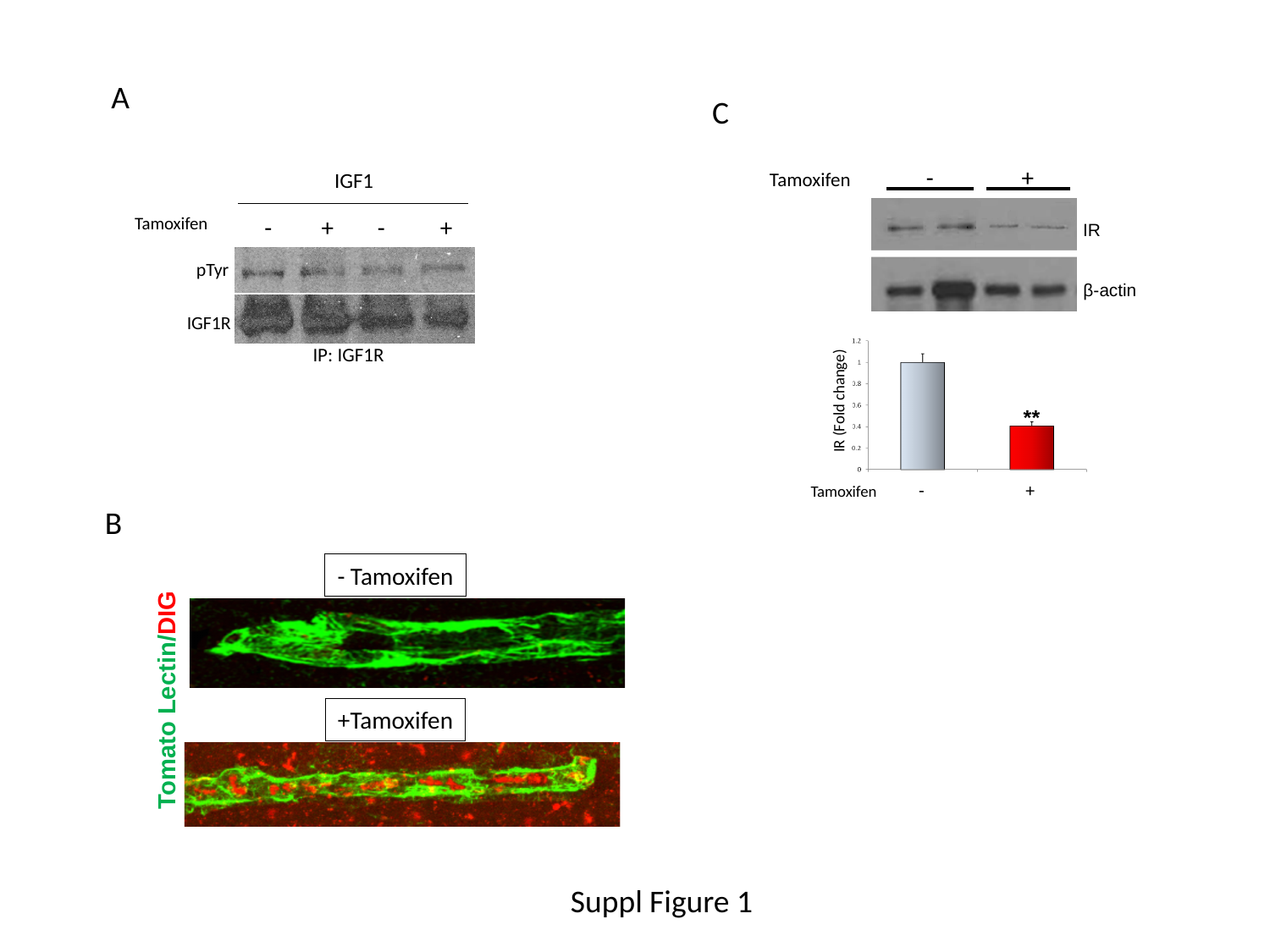

A
C
Tamoxifen - +
IR
β-actin
IR (Fold change)
**
Tamoxifen - +
IGF1
 - + - +
Tamoxifen
 pTyr
IGF1R
IP: IGF1R
B
- Tamoxifen
Tomato Lectin/DIG
+Tamoxifen
Suppl Figure 1
